# Aging macaques bridge the translational gap in perivascular space biology

**DOI:** 10.64898/2026.01.14.699534

**Authors:** William A Liguore, Daniel L Schwartz, Juan Piantino, Avanthika Rajendran, Greg Jensen, Joshua A Karpf, Mykyta M. Chernov, Randall L. Woltjer, Steven G Kohama, Lisa C Silbert, Alison R Weiss

## Abstract

The prevalence of MR-visible perivascular spaces (PVS) is age-dependent in rhesus macaques. Automated quantification in 94 male and female animals ranging 5-28 years of age demonstrated a robust association between age and PVS burden, with an anatomical distribution paralleling that of humans. Preliminary *ex vivo* validation confirmed that the MRI-detected tubular structures correspond to perivascular spaces. These findings establish the rhesus macaque as a tractable model for understanding the role of perivascular dysfunction in age-related brain vulnerability.

## MAIN

Perivascular spaces (PVS), or Virchow-Robin spaces, are fluid-filled channels bordered by astrocytic endfeet. They are typically microscopic but become visible with Magnetic Resonance Imaging (MRI) when large^1^, and have been posited as a structural signature of impaired solute clearance^1^, or “glymphatic” function. In humans, MR-visible PVS are typically elongated and cylindrical in shape, observed in white matter and basal ganglia structures, and isointense to cerebrospinal fluid (CSF) on MRI^2^. MR-visible PVS are associated with cerebral small vessel disease, neuroinflammation, and diffuse white matter hyperintensities^2-8^, and often appear before clinical symptoms^3,7-10^, underscoring their relevance as a biomarker of early-stage pathology. Aquaporin-4 (AQP4), a water channel protein on the astrocytic endfoot, is central to PVS function, and manipulations in mouse models have demonstrated its key role in regulating both CSF flow and amyloid beta (Aβ) clearance^11-14^. However, a fundamental limitation of common mouse models is that they do not develop MR-visible PVS. This creates a pressing need for a translational animal model capable of bridging the gap between mechanistic murine studies and human clinical research.

The macaque model of normative brain aging shares many features in common with humans including cognitive impairment, gray matter atrophy, increased white matter diffusivity, the proliferation of activated astrocytes and microglia, and the spontaneous development of mild Aβ plaques^15-30^. Yet, despite the close correspondence to human aging, the presence of MR-visible PVS has never been systematically evaluated in any nonhuman primate (NHP) species. Here, we show that macaques also spontaneously develop MR-visible PVS that increase in prevalence with advancing age, providing the first evidence that this key clinical phenotype is present in a translational NHP model.

Standard practice to assess PVS on MRI has historically involved manual identification by trained observers on one, or a handful, of axial slices. However, visual scoring methods offer only coarse sensitivity because they hinge on subjective field selection, omit volumetric or morphological information, are often inconsistent across studies, and are time-intensive to apply ^2^. More recently, high-throughput computational methods have been developed allowing automatic identification of perivascular spaces that can be applied brain-wide to identify and segment PVS on human clinical-grade MRI data^2^. One such approach has already been validated against manual axial ratings in human samples and is effective using T_2_-weighted (T2w) MRI contrast^2,31^.

To determine whether MR-visible PVS occur in aging macaques, we analyzed high-resolution whole brain T2w SPACE scans^32^ from 94 rhesus macaques (65F/29M; 5–28 years) (**Figure 1A**). Three 0.5mm isotropic T2w volumes per animal were acquired and averaged after co-registration to improve SNR. Images were aligned to the ONPRC18 T2w template^33^ and white matter masks in native space were generated. MR-visible PVS were identified using the MR imaging-based autoidentification of perivascular spaces (MAPS) algorithm^2,31^, which applies intensity-based clustering and morphology-guided filtering to segment tubular PVS-like structures (**Figure 1B**) and extract whole-brain counts and volumes (mm^3^).

**Figure 1.**
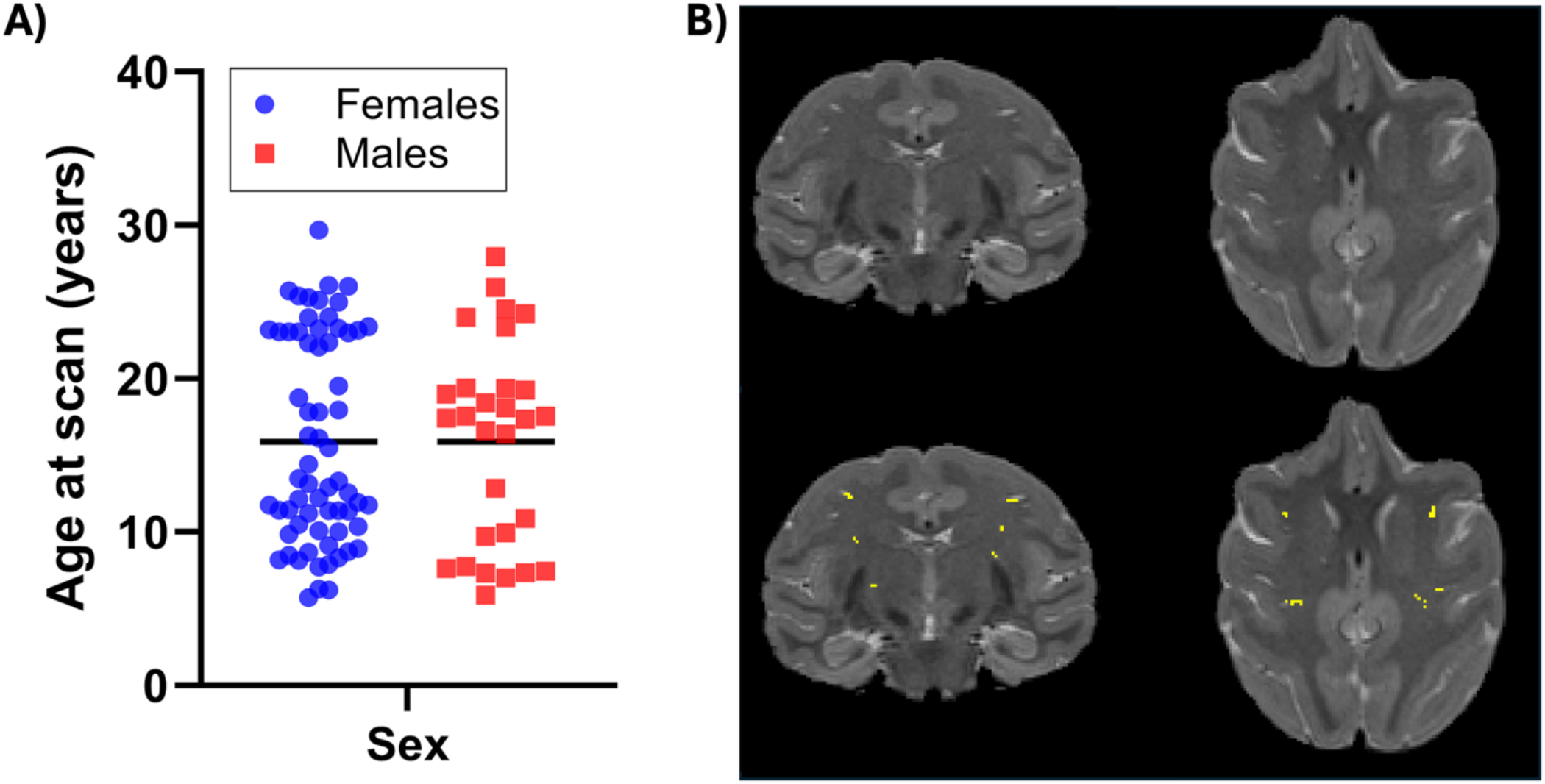
Demographics of study population and example *in vivo* MRI scans with automatic segmentation results. **(A)** Cohort of rhesus macaques studied includes n=65 females and n=29 males, ages 5-28 years old. Mean age for males was 15.87 years, while females had a mean age of 15.88 years. **(B)** Top row shows example coronal and axial T_2_-weighted MRI from a single animal, bottom row illustrates the results of MAPS PVS segmentation (yellow clusters).

MR-visible PVS were robustly detectable across the cohort and burden was age dependent. Both total PVS count (R(92)=0.5717, p<0.0001) and total PVS volume (mm^3^) (R(92)=0.5070, p<0.0001) positively correlated with age (years) (**Figure 2A,B)**, and these relationships strengthened when normalized by individual white-matter volume (mm^3^) (normalized count: R(92)=0.6778, p<0.0001; normalized volume: R(92)=0.6085, p<0.0001) (**Figure 2C,D)**. Linear regression analyses further quantified the strength of these age associations. Total PVS count and volume increased significantly with age (count: β = 1.134, R^2^ = 0.3268, F(1,92)=44.67, p<0.0001; volume: β = 0.9134, R^2^ = 0.2570, F(1,92)=31.82, p<0.0001). Normalization to white-matter volume yielded even steeper age-dependent trajectories (normalized count: β = 0.0001922, R^2^ = 0.4594, F(1,92)=78.18, p<0.0001; normalized volume: β = 0.00015, R^2^ = 0.3703, F(1,92)=54.10, p<0.0001). No significant sex differences were observed. A frequency heatmap revealed that the centrum semiovale had the highest incidence of MR-visible PVS (**Figure 2E)**, paralleling the distribution of human subjects’ (**Figure 2F)**^34-36^. Automated estimates closely matched expert blinded evaluation: MAPS counts strongly correlated with single-slice counts in n=59 subjects (R(57)=0.737, p<0.0001), and voxel-level inspection in n=56 subjects confirmed near-perfect correspondence between MAPS detections and observer-verified clusters (R(54)=0.995, p<0.0001).

**Figure 2.**
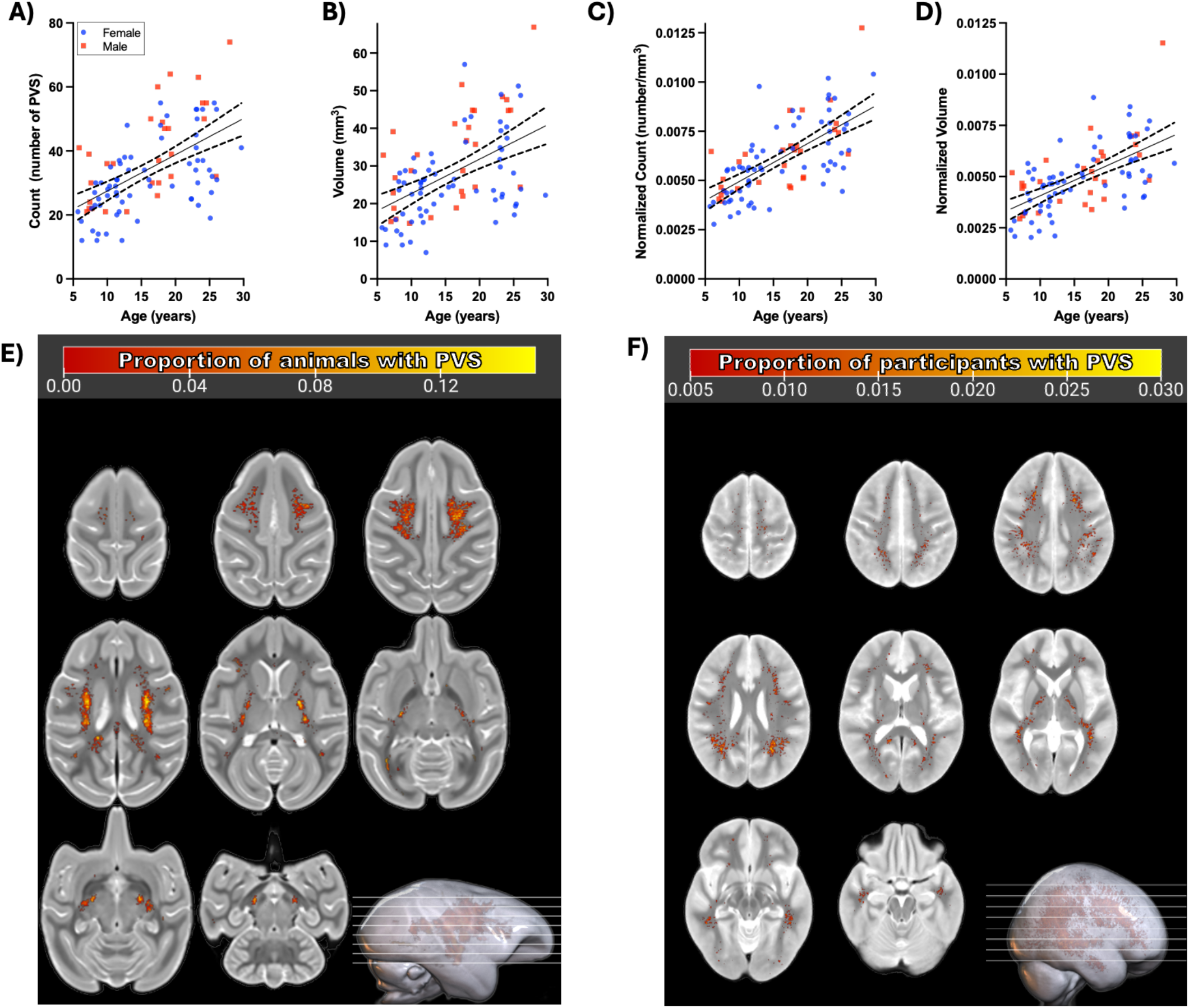
Whole brain assessment of MR-visible PVS. **(A)** Scatterplot showing the relationship of brain wide MR-visible PVS count, and **(B)** total brain wide PVS volume (mm^3^) with age; **(C, D)** shows the same data normalized to the volume of the white matter search area. **(E)** Heatmap of all n=94 individual PVS masks warped into a common space overlaid on ONPRC18 NHP brain template. Color overlay corresponds to the proportion of animals in the dataset with PVS identified in that location, and reveals that areas of the centrum semiovale and internal capsule have the highest incidence of MR-visible PVS. **(F)** A proportion heatmap of n=118 human participant’s PVS masks warped into common space and overlaid onto a study specific template that was previously reported^34^ is provided for spatial comparison to human regional white matter PVS distribution.

To corroborate the anatomical substrate of the MRI-defined structures, we conducted a qualitative *ex vivo* validation in a single aged macaque (26y) that was included in the *in vivo* dataset. The central goal of this case-study was to provide preliminary anatomical evaluation of a single radiological feature. To accomplish this, 0.25mm isotropic post-mortem MRI (**Figure 3A**) was collected to facilitate direct spatial alignment between MRI and matched histological sections (**Figure 3B**). In the feature investigated, trichrome staining showed a distinct separation between the vessel wall and surrounding parenchyma, together with collagen deposition, both hallmarks of enlarged PVS^6^ (**Figure 3C,D**). Additional stains of consecutive tissue sections provided further complementary, although not exhaustive, characterization of the structure and surrounding tissue. Specifically, smooth muscle actin (SMA) labeling marked a smooth-muscle containing vessel (**Figure 3E**). Although we did not undertake exhaustive vessel-type classification across multiple sections, these results are consistent with the observation that both arterial and venous segments can form MR-visible PVS^37^. Cresyl violet staining confirmed preserved cytoarchitecture surrounding the PVS (**Figure 3F**). Glial fibrillary acidic protein (GFAP) staining highlighted astrocytic processes bordering the space, and ionized calcium-binding adapter molecule 1 (IBA1) positive staining identified microglia within the parenchyma as well as cells within the PVS. Aquaporin 4 (AQP4) immunoreactivity was observed along astrocytic endfeet adjacent to the vessel wall (**Figure 3G–I**). Together, these data support the identity of this histological feature as a PVS^3^. While more in-depth follow-up histological studies are warranted, data from this single-subject qualitative survey suggest that the tubular hyperintensities detected *in vivo* correspond to bona fide perivascular channels with conserved cellular and extracellular features, and provide biological grounding for the MRI phenotype reported here.

**Figure 3.**
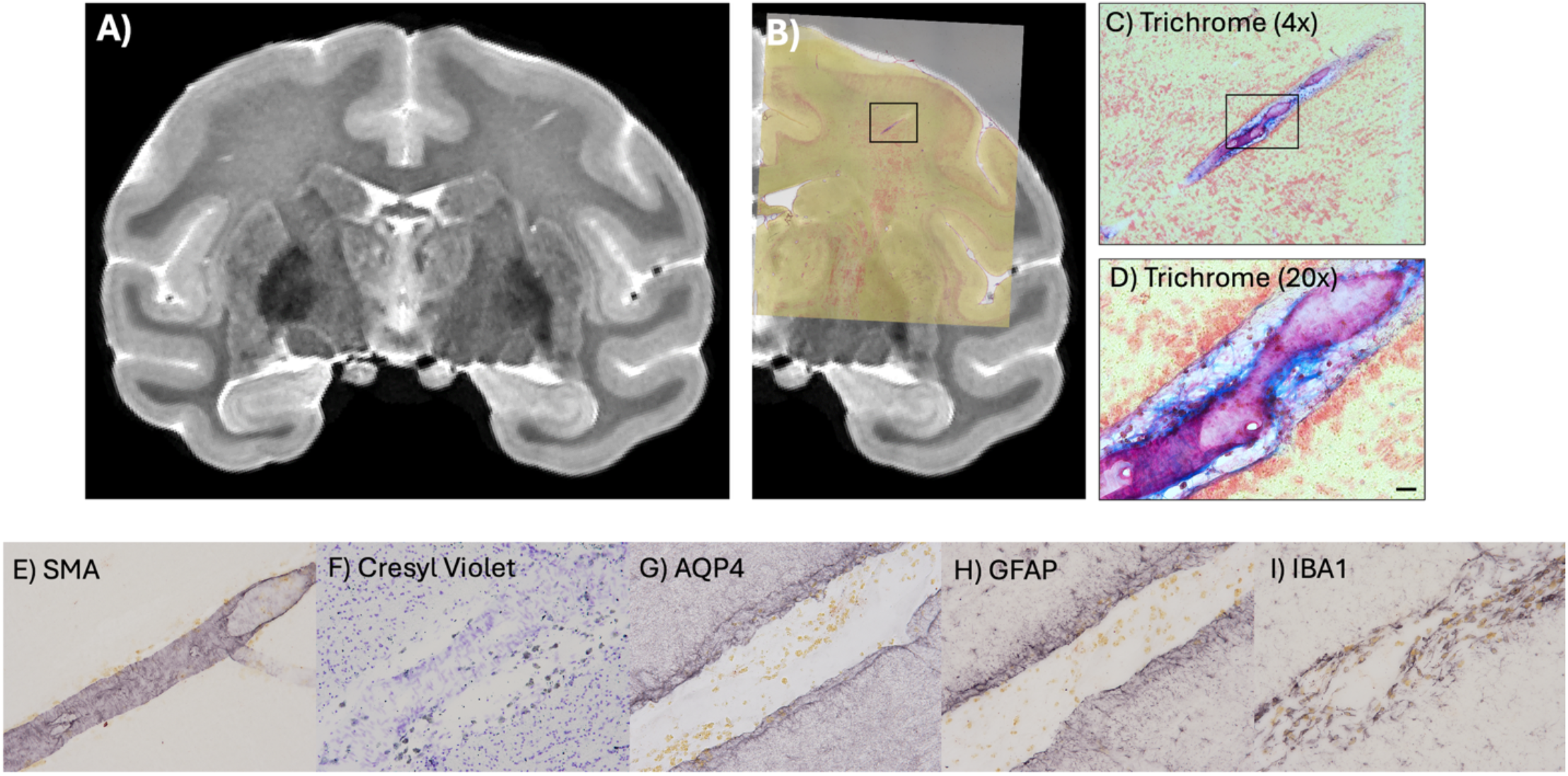
Ex vivo anatomical characterization of an MR-visible PVS. **(A)** Post-mortem T2-weighted MRI with 0.25mm isotropic resolution collected from a female macaque aged 26 years. **(B)** Histological staining (Masson’s trichrome) with post-mortem MRI co-alignment. Black inset box highlights the specific structure examined. **(C)** Image of the inset from (B) shown at 4x magnification. **(D)** Image of the black inset box from (C) shown at 20x magnification, with a scale bar in bottom right of 50um. Masson’s trichrome labels nuclei (dark blue), collagen (blue), and smooth muscle (red). Pannels **(E-I)** illustrate consecutive 40um thick tissue sections of the same structure, all captured at 20x magnification, with the following stains: **(E)** smooth muscle actin (SMA) labels arteries, **(F)** cresyl violet labels Nissl bodies, **(G)** aquaporin 4 (AQP4) labels astrocytic endfeet, **(H)** glial fibrillary acidic protein (GFAP) labels astrocyte cytoskeleton, and **(I)** ionized calcium-binding adapter molecule 1 (IBA1) labels microglia and macrophages.

These results demonstrate, for the first time, that rhesus macaques naturally develop MR-visible enlarged PVS as they age, thereby establishing the first naturally occurring NHP model of this key radiological marker. The strong correspondence between automated MAPS quantification and expert evaluation confirms that this phenotype can be measured with high reliability in macaques, enabling rigorous whole-brain quantification in large cohorts. Our findings also mirror several defining features of the human phenotype: age-associated increases^36,38^, predominance within deep and subcortical white matter^31,35,36,38^, and morphological characteristics consistent with tubular perivascular channels^31^. High-resolution fixed-tissue imaging captured the same tubular structure identified *in vivo*, and the alignment of MRI with microscopy data revealed histological features characteristic of a perivascular channel analogous to that of humans^3,6^. This cross-modal correspondence strengthens confidence that the *in vivo* MR-visible structures identified here are indeed perivascular spaces, rather than nonspecific white-matter abnormalities.

The availability of a naturally occurring NHP model for enlarged PVS creates new opportunities for mechanistic and translational research. The mechanisms underlying PVS enlargement are poorly understood, with hypotheses ranging from reduced AQP4 expression^39^, sleep fragmentation^40^, atrophy of surrounding tissues^41^, to altered cardiovascular pulsatility^42,43^. Rodent models have been instrumental in defining fundamental glymphatic mechanisms^11-14^; however, they largely fail to replicate the human radiological phenotype. Beyond one rat model of cardiovascular disease displaying hippocampal PVS^44^, no murine models have been reported to spontaneously develop MR-visible PVS with the white-matter dominance seen in humans. Furthermore, because rodent research primarily focuses on the lissencephalic cortical mantle, it fails to address the perivascular biology of deep white matter, the specific compartment where enlargement is most clinically relevant in the aging human brain. Macaques, by contrast, share key features of human aging^15-29^ and develop MR-visible PVS in white matter, providing an integrative model in which perivascular dysfunction can be studied alongside other age-related processes. Future longitudinal studies in this species will allow investigators to determine when PVS enlargement emerges relative to other neuropathological milestones, what vascular and tissue characteristics might precede enlargement, and how vascular or inflammatory perturbations modulate PVS trajectories. More importantly, this model enables controlled experimental manipulation of candidate mechanisms underlying PVS enlargement. Age-related changes in vascular stiffness, AQP4 depolarization, sleep fragmentation, cerebrovascular inflammation, and glymphatic flow can be measured and modulated in macaques using behavioral, pharmacologic, surgical, and imaging approaches unachievable in human clinical studies. The capacity to track global PVS burden and individual PVS morphology before and after such interventions in controlled experimental settings provides a powerful new framework for determining causal contributors to perivascular dysfunction and for evaluating therapeutic interventions targeting clearance pathways.

By revealing that MR-visible enlarged PVS arise spontaneously in aging rhesus macaques, and by validating their radiological and histological signatures, this work establishes the NHP platform for mechanistic, longitudinal, and interventional studies of the perivascular system in aging. This model will enable deeper understanding of how perivascular dysfunction contributes to late-life brain vulnerability and will offer a critical foundation for the development of strategies aimed at preserving perivascular health in aging and disease.

## ETHICS STATEMENT

All procedures received approval from the Oregon National Primate Research Center (ONPRC) Institutional Animal Care and Use Committee (IACUC) and were performed within the ONPRC animal care program. This program maintains full AAALAC International accreditation and operates in strict accordance with the *Guide for the Care and Use of Laboratory Animals* (8th edition), the Animal Welfare Act, and the Public Health Service Policy on Humane Care and Use of Laboratory Animals.

## ACKNLOWEDGEMENTS

We thank the staff of the Oregon National Primate Research Center (ONPRC) for expert animal care, veterinary support, and assistance with sample collection. We are grateful to the staff of the ONPRC Primate Multimodal Imaging Center (PMIC), the OHSU Advanced Imaging Research Center (AIRC), and the Oregon Alzheimer’s Disease Research Center Neuroimaging and Pathology Cores for guidance and technical support. Services reported in this publication provided by the NHP Aging Resource were supported by the National Institute on Aging and the Office of the Director, National Institutes of Health.

## DATA AVAILABILITY

The data that support the findings of this study are available from the corresponding author upon reasonable request and establishment of appropriate Material Transfer Agreements.

## CRediT STATEMENT

**William A Liguore:** data curation, formal analysis, investigation, project administration, visualization, writing – original draft, writing – review & editing; **Daniel L Schwartz:** formal analysis, methodology, software, writing – review & editing; **Juan Piantino:** validation, writing – review & editing; **Avanthika Rajendran:** validation, writing – original draft; **Greg Jensen:** formal analysis, supervision; **Joshua A Karpf:** methodology, writing – review & editing; **Mykyta M. Chernov:** conceptualization, methodology, writing – review & editing; **Randall L. Woltjer:** conceptualization, writing – review & editing; **Steven G Kohama:** funding acquisition, resources, writing – review & editing; **Lisa C Silbert:** funding acquisition, methodology, resources, writing – review & editing; **Alison R Weiss:** conceptualization, formal analyses, funding acquisition, investigation, methodology, supervision, writing – original draft, writing – review & editing.

## ONLINE METHODS

### Demographics

We assessed MR-visible PVS in a cohort of 94 rhesus macaques (65F/29M) spanning an age range representing early adulthood to advanced old age (5-28 years). All animals were housed at the ONPRC with ad libitum access to water, two daily chow rations supplemented with fresh produce, and 12HR light/dark cycle.

### *In vivo* MRI Scan Acquisition

Three T_2_-weighted sampling perfection with application optimized contrasts using different flip angle evolution (SPACE)^32^ scans were acquired with 0.5mm isotropic voxels (TE/TR = 385/3200 ms, flip angle = 120°, 320 x 320 x 224) for each animal in this study (total acquisition time 29 minutes 42 seconds). Scans were acquired on a Siemens Prisma whole-body 3 Tesla (T) MRI instrument utilizing a 16 channel pediatric head RF coil. The three scans were subsequently co-aligned and averaged to improve SNR. The merged whole head image was aligned to the ONPRC18 T_2_-weighted whole head template^33^ using b-spline nonlinear transformations in ANTs^45^, and the resulting inverse-warp field applied to generate a brain mask in native space. This was used to skull strip the merged image. The skull-stripped brain was then intensity bias corrected using the ‘N4BiasFieldCorrection’ tool in ANTs to account for field inhomogeneity^46^. A second ANTs registration step was then run to align the skull-stripped brain image to the ONPRC18 T_2_-weighted brain template using b-spline nonlinear transformations. The resulting warp fields were then used to generate white matter masks (WM) in native space.

### PVS Segmentation

We applied an automatic detection method to segment PVS^2,31^. Briefly, this program utilized the distribution of local differences in image intensity to identify clusters of voxels in the white matter mask which are hyperintense relative to nearby tissue. The 3D voxel clusters were then modelled as individual objects and constraints on linearity, width, and size were applied (MATLAB 2019A). The clusters which survived the constraints were collated in a 3D image and statistical dataset.

### Statistical Analysis

We calculated the number (count) and volume (mm^3^) of PVS clusters for each scan. These values were also normalized by dividing by each animal’s WM mask volume to account for differences in brain size between individuals. Two-tailed Pearson correlations and linear regressions were used to assess the relationships between each measure and age and sex. All statistical analysis was performed in GraphPad Prism (v.10). To provide spatial context to the findings, the binary PVS masks were co-aligned to the ONPRC18 template^33^ (ANTs), summed, and then divided by the total number of animals in the dataset to generate a heatmap showing the proportion of population with PVS identified in each voxel throughout the brain. A lightbox image of the population frequency heatmap was generated in MRIcro GL (www.nitrc.org).

### Post-mortem tissue preparation

The brain of one subject (26y F) with a high PVS burden was collected for a follow-up histological case-study. The animal was sedated with 10mg/kg ketamine and subsequently a lethal dose of pentobarbital was administered. After deep anesthesia was verified by lack of palpebral reflex, an incision was made opening the thoracic cavity. The animal was exsanguinated, then the cerebrovasculature was flushed by perfusing 1 liter of normal saline through the common carotid artery. Following saline, 1 liter of 4% paraformaldehyde (PFA) in 0.1M phosphate buffer was flushed through the vasculature. The brain was then extracted from the skull and further fixed in 4% PFA for 48hr.

### Post-mortem MRI scan acquisition

The paraformaldehyde (PFA) fixed whole brain was submerged in Fluorinert™ (FC-770, 3M) in a plastic container and scanned in the Siemens 3T MRI with an 8-channel receive only head coil (Rapid Biomedical GmbH). A T_2_-weighted SPACE experiment with 0.25mm isotropic voxels (TE/TR=310/3000, 80mm x 320 x 320) was collected. Following this, the fixed whole brain was sectioned coronally into 4mm thick slabs using a rhesus brain matrix (ASI Instruments, Inc.). To facilitate alignment with microscopy images, the 4mm thick brain slabs were assembled into approximately anatomically correct positions in a plastic jar separated by acrylic spacers aligned with 3 nylon rods and 2mm thick washers. This jar was filled with Fluorinert™ and scanned in the Siemens 3T MRI utilizing the pediatric head coil previously described. Three repetitions of T_2_-weighted SPACE were collected with the same parameters as the in-vivo scans. After averaging, the individual slabs were manually extracted and reassembled as a whole brain image using FSL and FIJI tools^47^.

### Neuropathology

After scanning, coronal slabs were transferred to a 30% sucrose solution for 1 week and then sectioned on a freezing microtome at 40µm thick and stored in a cryoprotectant solution (30% sucrose, 30% ethylene glycol, 0.2M phosphate buffered saline). Consecutive 40µm thick tissue sections were washed to remove cryoprotectant solution, endogenous peroxidases were quenched with 0.25% sodium meta-periodate (Sigma), and non-specific antibody binding was blocked with 5% normal goat serum (Gibco). Tissue sections were incubated in primary antibody solution overnight at room temperature. Antibodies included glial fibrillary acidic protein (1:2000, Dako), ionized calcium-binding adaptor molecule 1 (1:1000, WAKO), smooth muscle actin (1:500, Dako), and aquaporin-4 (1:500, Millipore). A one-hour incubation in secondary antibodies (1:500, Jackson Labs), and Avidin Biotin Complex solution (Vector Labs) preceded development in 0.5% 2’-4’ diaminobenzidine (Sigma) with 2.5% nickel II sulphate (Sigma). A series of 40µmthick tissue sections were also stained using Masson’s Trichrome kit (Newcomer Supply, Inc.) and cresyl violet (1%, Sigma). All sections were imaged on a BX51 Olympus microscope.

